# An embryonic stem cell-specific heterochromatin state allows core histone exchange in the absence of DNA accessibility

**DOI:** 10.1101/2020.05.22.110742

**Authors:** Carmen Navarro, Simon J Elsässer

## Abstract

Nucleosome turnover concomitant with incorporation of the replication-independent histone variant H3.3 is a hallmark of regulatory regions in the animal genome. In our current understanding, nucleosome turnover is universally linked to DNA accessibility and histone acetylation. In mouse embryonic stem cells, H3.3 is also highly enriched at interstitial heterochromatin, most prominently intracisternal-A particle endogenous retroviral elements. Interstitial heterochromatin is established over confined domains by the TRIM28/SETDB1 corepressor complex and has stereotypical features of repressive chromatin, such as H3K9me3 and recruitment of all HP1 isoforms. Here, we demonstrate that fast histone turnover and H3.3 incorporation is compatible with these hallmarks of heterochromatin. Further, we find that histone H3.3 is required to maintain minimal DNA accessibility in this surprisingly dynamic heterochromatin state. Loss of H3.3 in mouse embryonic stem cells elicits a highly specific opening of interstitial heterochromatin with minimal effects on other silent or active regions of the genome.

## INTRODUCTION

Histone H3.3 is an evolutionary conserved variant of the canonical histone H3 proteins, termed H3.1 and H3.2 in mammalian cells ^1^. First described to associate with regions of active transcription in *Drosophila* ^2^, H3.3 has been recognized as the sole substrate for replication-independent chromatin assembly in mammalian cells ^3^. While H3.1/2 expression and replication-dependent incorporation is limited to S phase, H3.3 is expressed through the cell cycle and incorporated at sites of dynamic nucleosome turnover. A multitude of genome-wide studies have identified such dynamic chromatin at promoters, enhancers, gene bodies and origins of replication ^4–13^. In these instances, nucleosome turnover is thought to be a consequence of chromatin remodeling, transcriptional activity, nucleosome-destabilizing DNA sequences, competition with other chromatin binding factors, or a combination thereof. While histone H3.3 itself has been proposed to destabilize the nucleosome, it is unclear if H3.3 has a causal role in promoting nucleosome turnover ^14–16^. Histone H3.3 has been shown to be enriched in histone posttranslational modifications (PTMs) characteristic of euchromatin, i.e. H3K4me3, H3K9ac, H3K27ac ^17,18^. Accumulation of active PTMs on histone H3.3 is sufficiently explained by their co-occurrence at sites of active enhancers or transcription, and H3.3 appears not to be required for maintaining those active PTMs. However, a notable exception represents phosphorylation of Ser31, a H3.3-specific residue. H3.3Ser31 phosphorylation facilitates rapid activation of genes ^19,20^, thus providing a mechanistic link between H3.3 incorporation and gene activation. In summary, histone H3.3 is a well-established component of euchromatin, acting as a replacement histone for a range of dynamic processes. While H3.3 has to be considered predominantly neutral to the underlying dynamic process, it can – as exemplified in the specific instance above – partake mechanistically in establishing open chromatin.

In mouse embryonic stem cells (ESC), a considerable fraction of H3.3-enriched regions do not fall into the active regions outlined above, but colocalizes with histone H3 Lys 9 trimethylation (H3K9me3) modification at interstitial heterochromatin ^21^. Re-ChIP experiments suggested that H3.3 and H3K9me3 coincide in the same nucleosome. Interstitial heterochromatin in mouse ESC spreads over relatively small genomic distance (∼10kb), is established by a Tripartite motif-containing 28 (TRIM28 also known as KAP1, TIF1b) co-represser/SETDB1 histone methyltransferase complex and includes a subset of endogenous retroviral elements (ERVs), most prominently Intracisternal A-type particle (IAP), and imprinted genes ^22–27^. KAP1 is recruited to foreign DNA elements through numerous DNA sequence specific Krüppel-associated box containing Zing-Finger proteins (KRAB-ZFPs).

Interstitial heterochromatin is highly CpG-methylated, enriched in linker histone and bound by HP1 family proteins, the stereotypical H3K9me3 readers ^28–30^. HP1 proteins have been shown to bridge nucleosomes and compact chromatin into phase-separated liquid condensates ^31,32^. HP1 and linker histone render DNA in an inaccessible state ^32^, and underlying genes/repeats are refractory to activating factors ^33^.

Enrichment of H3.3 at H3K9me3 heterochromatin regions, so far uniquely observed in mouse ESC, poses a conundrum: do these regions show fast nucleosome turnover as known for euchromatic H3.3-enriched regions? Does the presence of H3.3 coincide with DNA accessibility, and how would such dynamic properties be reconciled with a silent, condensed heterochromatin structure? Here, we reveal surprisingly dynamic properties of interstitial heterochromatin in ESC and find well-known hallmarks of heterochromatin are compatible with rapid exchange of core histones while maintaining DNA in an inaccessible state.

## RESULTS

### H3.3 is a pluripotent stem cell-specific component of interstitial heterochromatin

Several studies have assessed genome-wide H3.3 dynamics in ESC, either measuring incorporation or turnover of ectopically tagged H3.3 ^5,8,9,34^, albeit exclusively looking at dynamics related to enhancers and transcription. A recent study assessed nucleosome turnover using a method termed ‘time-ChIP’ in mouse embryonic stem cells as well as neural stem cells (NSC) ^9^, reporting high nucleosome turnover at enhancers correlating with DNA accessibility. For the time-ChIP method, SNAP-tagged H3.3 was expressed from a Tet-controlled transgene and biotin-labeled using the SNAP-tag at the onset of a time course (0h). The biotin-labeled fraction then was observed over a time range of 3-12 hours (Figure 1a, b). Dynamic regions are marked by H3.3 at 0h and gradually decay towards the genome-wide average, implying replication-independent turnover of H3.3 ^9^.

**Figure 1.**
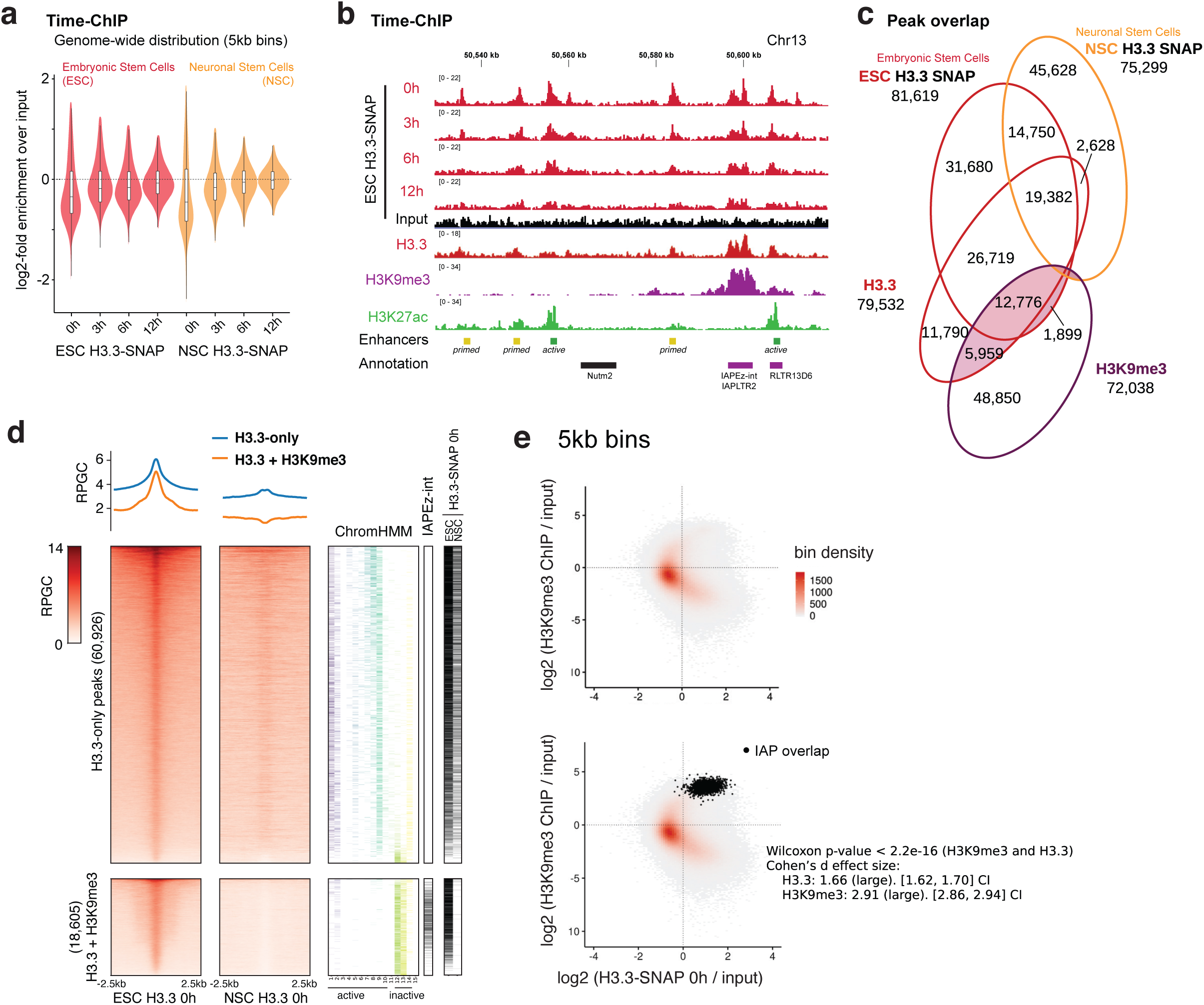
Genome-wide distribution of ectopic H3.3-SNAP in mouse ESC and NSC shows ESC-specific enrichment at interstitial heterochromatin. **a** Violin plots showing genome-wide distribution of biotin-labeled H3.3-SNAP ^9^ in 5kb windows, in mouse embryonic stem cells (ESC) and neuronal stem cells (NSC). Biotin labeling of H3.3-SNAP was performed at 0h and followed over subsequent 3, 6 and 12h ^9^. Each datapoint is calculated as the log2 value of the mean coverage in each sample bin divided by the mean coverage value of its corresponding input sample bin. **b** Genome tracks showing example region of time-ChIP ^9^ in ESC showing overlap of H3.3-SNAP and endogenous H3.3 ^21^ with H3K27ac ^20^ at enhancers and H3K9me3 ^51^ at IAP ERVs. All tracks were normalized to Reads Per Genomic Content (RPGC). See also Extended Data Figure 1 for controls and additional tracks. **c** Venn diagrams showing overlap of H3.3-SNAP 0h ^9^ peaks with endogenous H3.3 peaks in ESC ^21^, H3K9me3 ^51^ and H3.3-SNAP 0h peaks in NSC ^9^. **d** Read density heatmaps and average profiles of H3.3-SNAP in ESC and NSC, at ESC H3.3 peaks ^21^. Peaks were separated in two categories “H3.3-only” and “H3.3+H3K9me3” according to coincidence with H3K9me3 peaks ^21^. Overlap with 15 State ChromHMM ^36^ states, IAP ERV elements (from repMasker) and H3.3-SNAP peaks in ESC and NSC is shown as additional heatmaps where the coincidence with a chromatin state or peak from the respective annotation set is indicated with a colored or black line. **e** Scatter plot showing relationship between H3.3 ^21^ and H3K9me3 ^51^ assessed over 5kb bins genome-wide. Color scale represents density of underlying data points (5kb bins). Bottom graph shows bins overlapping with IAP ERVs in black. Wilcoxon rank sum (p<2.2e^-16^) and Cohen’s effect size d (H3.3: 1.66+/-0.04; H3K9me3: 2.91+/-0.05) tests show significant enrichment for both at H3.3 and H3K9me3 in these 5kb bins. Overlap with additional repeat families are shown in Extended Data Figure 3.

We reanalyzed the time-ChIP datasets with respect to the known heterochromatic enrichment of H3.3. H3.3-enriched regions in ESC called from the initial time point (0h) overlapped well with those determined for endogenous histone H3.3 ^21^, including those peaks sharing H3.3 and H3K9me3 (H3.3+H3K9me3) (Figure 1c,d). As an example, H3.3 enrichment tracked well with known ESC enhancers and H3K27ac marks, but to similar levels with a nearby IAP ERV (Figure 1b, Extended Data Figure 1a). NSC showed globally similar turnover kinetics (Figure 1a, Extended Data Figure 1b, c) and maintained many of the H3.3 peaks defined in ESC (Figure 1c,d). However, among those H3.3 peaks specifically lost in NSC were essentially all H3.3+H3K9me3 peaks (Figure 1c,d), as exemplified by an IAP ERV localized in the Hist1 histone gene cluster (Extended Data Figure 1b). Thus, as previously concluded ^21^, co-enrichment of H3.3 and H3K9me3 is a feature of pluripotent stem cells which appears to be lost early upon differentiation. Interestingly, H3K9me3 and H3.3 has also been studied genome-wide during reprogramming of mouse fibroblasts to induced-pluripotent cells ^35^; while H3K9me3 becomes enriched over IAP ERV in pre-iPSC, H3.3 is absent at this stage (Extended Data Figure 1c). Pre-iPSC are stable late-stage reprogramming intermediates that can be converted to the pluripotent state through MEK inhibition. Thus, these data corroborate a link between pluripotency and the acquisition of H3.3 at interstitial heterochromatin, where H3K9me3 has already been established during reprogramming.

Annotating H3.3 peaks with a 15-state ChromatinHMM model generated with seven histone modifications and RNA Polymerase II (RNAP2) profiles ^36,37^, we found that H3.3-only peaks overlapped largely with active transcription (states 1-3) and enhancers (states 4-9) (Figure 1d, Extended Data Figure 2a). H3.3+H3K9me3 peaks were assigned to heterochromatin (states 12-14) (Figure 1d, Extended Data Figure 2a), defined by the absence of H3K4me3, H3K4me1, H3K27ac, H3K36me3 and RNAP2 ^37^, in line with the fact that H3K9me3 appears mutually exclusive with these signatures of active chromatin. Binning the genome into 5kb windows, we found a bimodal H3.3 distribution, with regions of very high and very low levels of H3K9me3 coinciding with the highest H3.3 enrichment (Figure 1e). Overlaying the 5kb bins with repeat annotation, we confirmed that the strongest H3.3+H3K9me3 co-enrichment was linked to the presence of an IAP element (Figure 1e, Extended Data Figure 3). Further, ETn and MusD elements also defined a subset of H3.3+H3K9me3 co-enriched regions (Extended Data Figure 3). It is known that a small fraction of ETn/MusD elements are active and, in fact, highly transcribed. This is evident from elongating RNA Polymerase II running into the 3’ flanking region of few elements, and the complete absence of H3K9me3 around these elements (Extended Data Figure 4a,b). Since we cannot unambiguously attribute ChIP-Seq reads within the highly conserved ETn/MusD internal sequence to specific instances, we are not able to discern how much H3.3 incorporation can be attributed to the small fraction of active versus majority of repressed copies. Therefore, we did not further focus on ETn/MusD elements. Amongst the IAP repeat family, none of the annotated instances showed similar evidence of transcriptional activity (Extended Data Figure 4c,d), and RNA-Seq transcript levels were 1-2 orders of magnitude lower than those of ETn (Extended Data Figure 4e).

### Rapid histone turnover at interstitial heterochromatin in the absence of DNA accessibility

Using the time-ChIP data, we next assessed the turnover of labeled histone H3.3 at H3.3-only and H3.3+H3K9me3 peaks. Decay of H3.3 towards background levels over the 12h chase period was observed at all peaks irrespective of their H3K9me3 status (Figure 2a). Turnover at H3.3+H3K9me3 enriched regions was also observed in another Tet-OFF dataset ^5^ (Extended Data Figure 5a). A different dataset, applying a pulse of tagged histone H3.3 ^34^, shows that newly synthesized histone H3.3 is incorporated rapidly at both H3.3-only and H3.3+H3K9me3 peaks (Figure 2b, Extended Data Figure 1a, 5). An orthogonal method to measure nucleosomal H3-H4 incorporation and turnover, CATCH-IT, relies on metabolic labeling of newly synthesized proteins with Azido-homoalanine (AHA) ^38^. CATCH-IT previously performed in mouse ESC ^6^ confirms the dynamic exchange of core histones at both H3.3-only and H3.3+H3K9me3 peaks (Figure 2c). Crucial for this method, all non-histone proteins and H2A/H2B dimers are stripped of chromatin biochemically before performing the ChIP, revealing the incorporation dynamics of newly synthesized H3-H4 units irrespective of the H3 variant. Thus, three independent datasets provide complementary evidence for nucleosome turnover at interstitial heterochromatin. While rapid H3.3 turnover has been attributed to ‘hyperdynamic’ promoter and enhancer nucleosomes ^8,9^, our analysis suggests that similar dynamics also apply to H3.3+H3K9me3 peaks.

**Figure 2.**
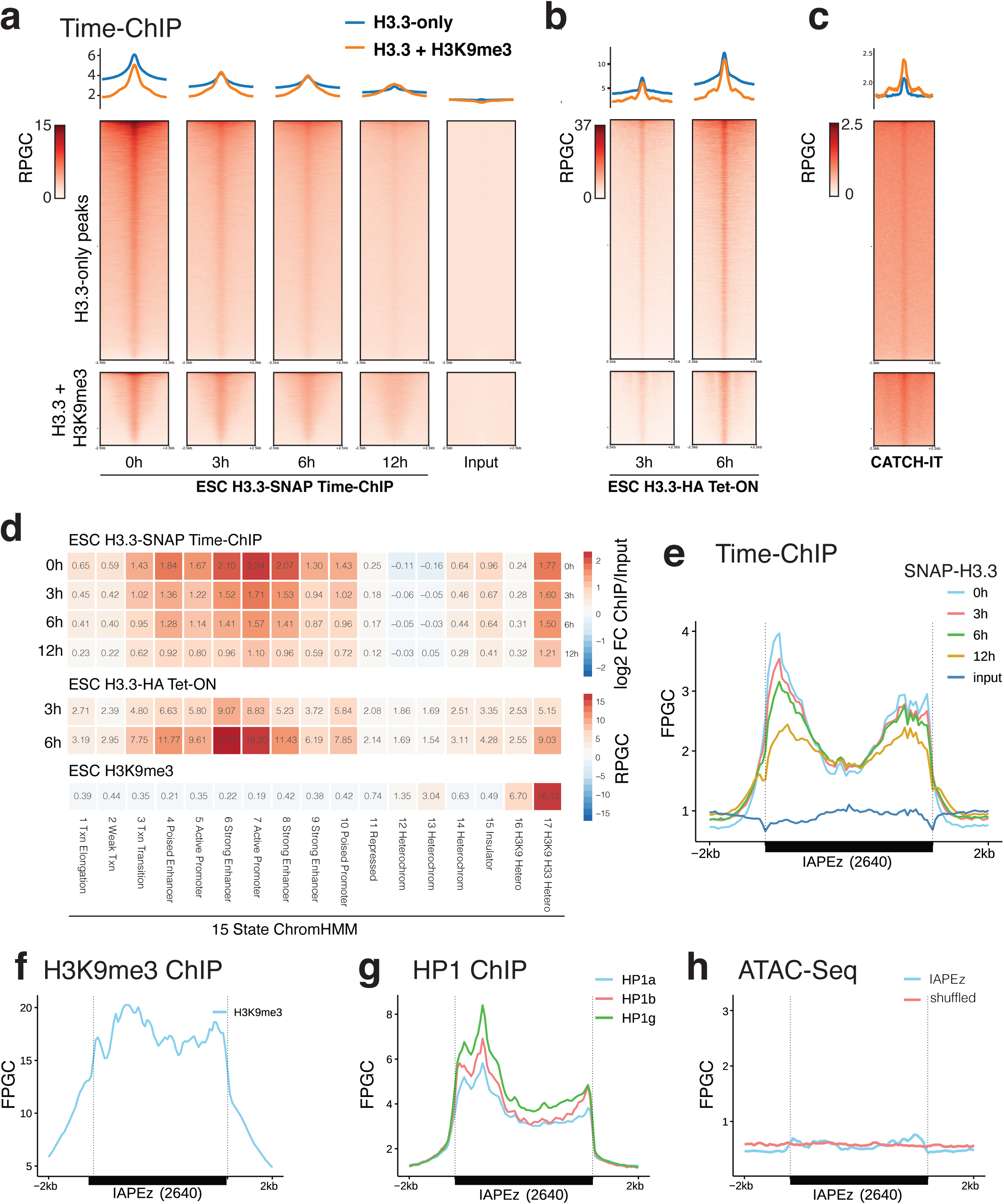
Histones are rapidly turned over at interstitial heterochromatin in the absence of DNA accessibility. **a**Read density heatmaps and average profiles of H3.3-SNAP time-ChIP data ^9^ over two classes of H3.3 peaks, H3.3-only and H3.3+H3K9me3 peaks. All data is normalized to Reads Per Genomic Content (RPGC) **b** Read density heatmaps and average profiles of H3.3-HA pulse data ^34^ over H3.3 peaks. **c** Read density heatmaps and average profiles of CATCH-IT data ^6^. **d** Mean read density heatmap showing enrichments of H3.3-SNAP time-ChIP (log2 fold-change over input) over 15 ChromHMM states ^37^ as well as H3K9me3 and H3.3+H3K9me3 enriched regions ^21^. Additional replicates are shown in Extended Data Figure 5b. For enrichment analysis by repeat families see also Extended Data Figure 6. **e** Average coverage of H3.3-SNAP time-ChIP over 2640 shared IAP ERVs. Fragments defined by paired-end reads were piled up and normalized by 1x Genome coverage (Fragment Per Genomic Content, FPGC). See Extended Data Figure 5c for coverage of uniquely mappable reads only. **f** Average coverage (FPGC) of H3K9me3 ChIP ^51^ 2640 shared IAP ERVs. **g** Average coverage (FPGC) of BioChIP for HP1 isoforms ^52^ over 2640 shared IAP ERVs. **h** Average profiles of DNA accessibility ^20^ over 2640 shared ERVs, and 2640 random (shuffled) genomic regions of matching size.

We next compared H3.3 dynamics across 15 ChromHMM states ^37^ and H3.3+H3K9me3 heterochromatic regions and found that highest histone H3.3 turnover is observed at active promoters and enhancers followed by H3.3+H3K9me3 regions (Figure 2d, ExtendedData Figure 5b). Analysing individual repeat families using UCSC RepeatMasker annotation, the ∼340bp IAPLTR1/1a long terminal repeats showed comparable turnover to active promoters and enhancers, followed by IAPLTR2 and internal IAP regions (Extended Data Figure 6). Examining H3.3 dynamics across full-length IAPLTR1 elements, it was interesting to note that histone turnover was not confined to individual positioned nucleosome but broadly occurred over LTRs and adjacent ∼2kb of internal region (Figure 2e). Remarkably, regions of highest turnover were also highly enriched for H3K9me3, concomitant with HP1 binding (Figure 2f-g). Prior analysis suggested that, as stereotypical for heterochromatin, DNA accessibility at ERVs is generally low ^29^, and analysis of recent ATAC-Seq data in mESC ^20^ confirmed that IAPLTRs neither show defined nucleosome-free regions nor broader domains of accessible DNA (Figure 2h). In summary, we find that histone turnover does not require or induce an increase in DNA accessibility at IAP ERVs (Figure 2f), Instead, it appears that histone H3.3-H4 is swapped into chromatin directly replacing an existing H3-H4 histone dimer or tetramer, hinting at the possibility of a concerted mechanism.

### Interstitial heterochromatin becomes accessible in the absence of histone H3.3

To achieve an exchange of core histones without rendering DNA at least transiently accessible, a tightly regulated supply of H3.3 would appear necessary. We thus wondered how a lack of H3.3 would affect DNA accessibility at H3.3+H3K9me3 regions. ATAC-Seq profiles of wildtype and H3.3 knockout cells have previously been compared ^20^. The authors note that DNA accessibility is largely unchanged across the genome in the absence of H3.3, including sites of rapid histone exchange, such as promoters and enhancers ^20^. Loss of H3.3 was thus fully compensated at promoters and enhancers, presumably by assembly or recycling pathways for H3.1/H3.2. In line with the original report, our re-analysis of the ATAC-Seq signals at H3.3-only peaks showed that DNA accessibility was not altered in H3.3KO cells (Figure 3a, b). Genome-wide analysis of 5kb bins and ChromHMM regions provided no evidence of systematic changes in DNA accessibility (Figure 3c, Extended Data Figure 7), suggesting that by large, loss of H3.3 had no effect on the packaging of the genome.

**Figure 3.**
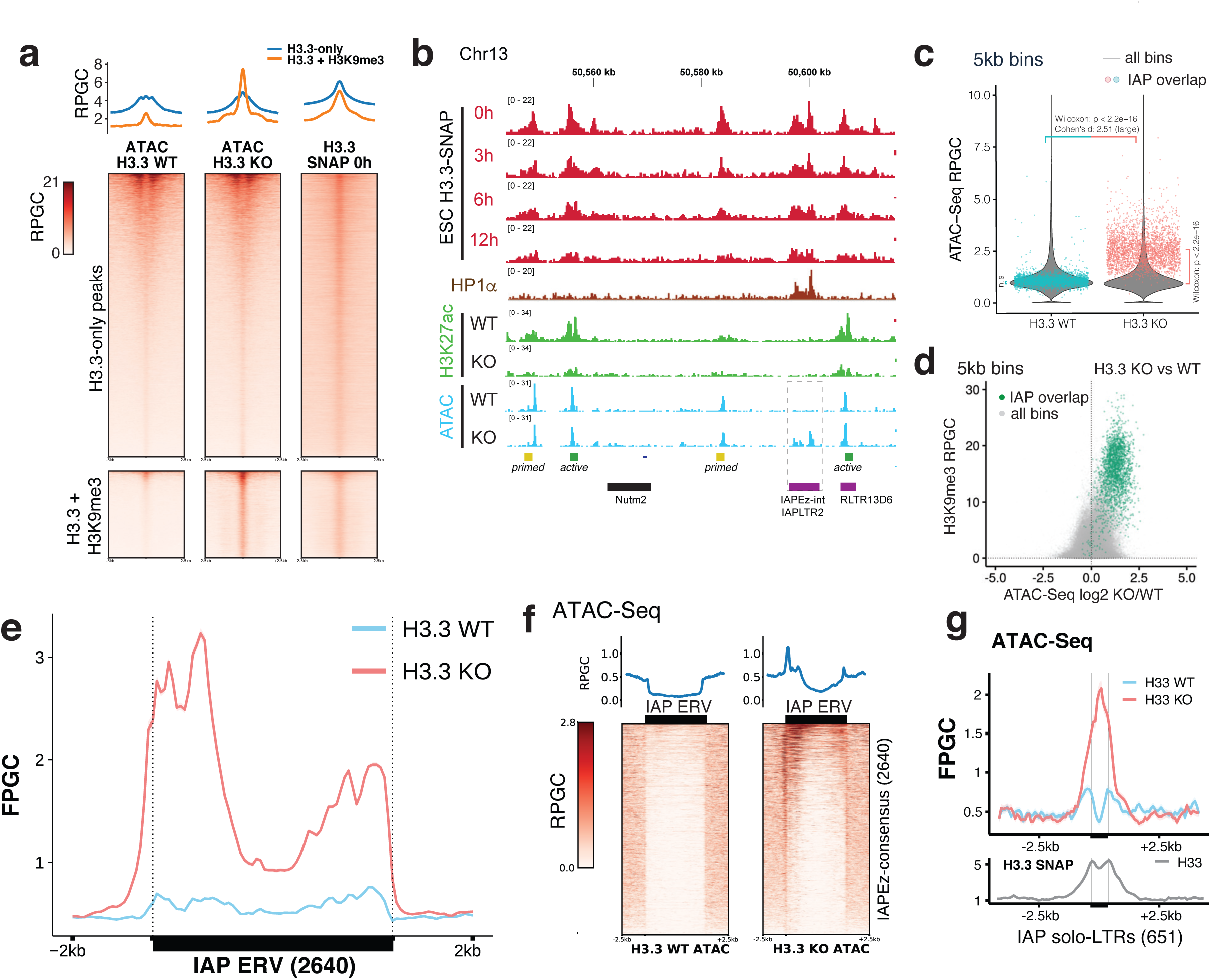
Interstitial heterochromatin becomes accessible in the absence of H3.3. **a**Read density heatmaps and average profiles of H3.3-SNAP time-ChIP ^9^ and ATAC-Seq ^20^ over two classes of H3.3 peaks, H3.3-only and H3.3+H3K9me3 peaks. See Extended Data Figures 7 and 8 for additional analysis **b** Genome tracks showing H3.3-SNAP time-ChIP ^9^, H3K27ac ^20^ and ATAC-Seq ^20^ signal over example region on Chr13. All tracks are normalized to Reads Per Genomic Content (RPGC). **c** Violin plots showing genome-wide distribution of ATAC-Seq signal in 5kb bins, in H3.3 wildtype (WT) and knockout (KO) cell line. Overlayed in color are 2797 individual bins overlapping with 2640 shared IAP ERVs. Wilcoxon rank test (p < 2.2e-16) and large Cohen’s effect size (d = 2.51+/-0.07) show significant increase in accessibility for IAP ERV containing bins in H3.3 KO. Genome-wide between H3.3 WT and KO shows a significant (p = 6.4e-14) difference but negligible effect size (d = 9.4e-6). IAP-overlapping bins are not different from genome-wide average in H3.3 WT (p = 0.97), but significantly different (p < 2.2e-16) in H3.3 KO. **d** Scatter plot of 5kb bins, showing H3K9me3 level versus log2-fold change in ATAC-Seq signal upon H3.3 knockout. Bins overlapping with IAP ERVs are drawn in green. **e** Average coverage of ATAC-Seq signal over 2640 shared IAP ERVs in H3.3 wildtype (WT) and knockout (KO) cell line. Fragments defined by paired-end reads were piled up and normalized to 1x Genome coverage (Fragment Per Genomic Content, FPGC). **f** Density heatmaps and average profiles of ATAC-Seq signal using only uniquely mappable reads, over 2640 shared IAP ERVs and flanking regions. See Extended Data Figure 7e for quantitative analysis and Extended Data Figure 9 for further comparison with H3.3 and H3K9me3 ChIP profiles **g** Average coverage (FPGC) of ATAC-Seq signal (top) and H3.3-SNAP ChIP using only uniquely mappable reads, over 651 IAP solo LTRs (mostly IAPLTR2).

On the background of this, it was striking to find a roughly threefold increase in DNA accessibility in H3.3KO cells specifically at H3.3+H3K9me3 peaks (Figure 3a, b). Genome-wide analysis of 5kb bins further confirmed in an unbiased manner that H3K9me3+H3.3 co-enrichment regions showed the highest fold-increase in DNA accessibility in the absence of H3.3 (Figure 3d, Extended Data Figure 7). This suggested that H3.3, while dispensable for proper packaging of other regions of the genome, is required for maintaining a closed chromatin state at H3.3+H3K9me3 heterochromatin.

As discussed above, a large fraction of H3.3+H3K9me3 heterochromatin is contributed by IAP ERVs (Figure 1e). Most IAP ERV containing 5kb bins moved from a low DNA accessibility (roughly genome-average) to the top quartile of accessible regions genome-wide in H3.3KO cells (Figure 3c). Although an increase in DNA accessibility could be indicative of a gain of euchromatic histone modifications, loss of H3.3 did not lead to a gain in histone H3K18, H3K27, H3K64 or H3K122 acetylation at H3.3+H3K9me3 peaks (Figure 3b, Extended Data Figure 8). This is in line with prior observations that loss of H3.3 leads to only a partial loss of H3K9me3 modifications and a moderate increase in transcriptional activity ^21,39^.

We next wondered if specific functional DNA elements of the IAP ERV would become accessible in the absence of H3.3. In contrast to the typically narrow peaks over promoters and enhancers corresponding to transcription factor binding sites, ATAC-Seq signal increased broadly across LTR and ∼2kb of internal region of IAP ERVs, in a profile that closely matched the distribution of H3.3 in wildtype cells (Figure 3e, see Figure 2e for comparison). Thus, H3.3 appears to play a specific role at IAP ERV in maintaining DNA in a tightly packaged and inaccessible state. Analysis of uniquely mappable paired-end reads within and adjacent to full-length IAP ERVs supports a widespread effect on many individual instances of the IAP ERV family (Figure 3f, Extended Data Figure 7e, Extended Data Figure 9). In addition to full-length or truncated IAP ERVs, the mouse genome contains many single LTRs (predominantly of the IAPLTR2 family). Using only uniquely mappable read pairs, we found that the 651 H3.3-enriched orphan LTRs also gained DNA accessibility in the absence of H3.3 (Figure 3g).

In summary, we find that interstitial heterochromatin maintains underlying DNA in an inaccessible state, a key characteristic of all heterochromatin. Despite this, interstitial heterochromatin is permissive to nucleosome turnover and histone H3.3 incorporation, providing a mechanistic explanation for the co-occurence of H3.3 and H3K9me3. According to the collective evidence from the data analyzed here, nucleosome eviction and assembly of new histone H3.3 nucleosomes must be accomplished through a mechanism that does not allow the underlying DNA to become accessible in the process. Loss of histone H3.3 may uncouple eviction of existing nucleosomes from immediate reassembly of a new nucleosome and consequently lead to the observed broad gain in DNA accessibility.

## DISCUSSION

In the current literature, heterochromatin is invariably associated with an inert chromatin packaging ^33^. Dynamic histone exchange is inherently viewed as a mechanism that opposes heterochromatin formation ^40^. Collectively, our analyses shed first light into a dynamic heterochromatin state that defies these stereotypical distinction of euchromatin and heterochromatin. While the purpose and mechanism remain only poorly understood, our study provides a novel hypothesis and functional insights for future mechanistic studies.

Histone H3.3 is known to be deposited at interstitial heterochromatin through the histone chaperone DAXX ^21,41–43^. DAXX has been proposed to contribute to ERV silencing through its histone chaperone activity and, independently, recruitment of histone deacetylases to ERVs ^21,39^. Loss of H3.3 also reduces global DAXX levels in mouse ESC, but H3.3+H3K9me3 regions did not gain histone H3 acetylation in the absence of H3.3 (Extended Data Figure 8). This suggests that impaired nucleosome assembly in the absence of H3.3 was the cause of the observed opening of chromatin (Figure 3a). The mechanism how existing nucleosomes are evicted to facilitate exchange with new H3.3 is unknown. The chromatin remodeler ATRX can form a complex with DAXX and has previously been linked to ERV silencing ^22,43–46^, yet it is unclear if ATRX contributes substantially to nucleosome turnover and H3.3 incorporation ^21,39,46^. The chromatin remodeler Smarcad1 has recently been linked to IAP ERVs in mouse ESC ^47^. Smarcad1 has further been shown to have nucleosomes sliding and eviction activity, thus representing another candidate for mediating H3.3 turnover at ERVs ^48,49^. Structural studies of the HP1-bound dinucleosome corroborate the view that heterochromatin can be accessible for nucleosome remodeling ^32^. We show that supply of H3.3 is necessary for maintaining chromatin structure at IAP ERVs, implying that existing nucleosome eviction is directly linked to DAXX-dependent deposition of new H3.3. A major question remains why ATP-dependent turnover of old and incorporation of new histone is required as part of a repressive chromatin domain. Transient eviction of nucleosomes could allow KRAB zinc-finger proteins to sample the DNA more effectively and thereby facilitate recruitment of TRIM28 corepressor complex. Alternative models could be proposed where nucleosome eviction represents a mere side effect of a more widespread chromatin remodeling activity in embryonic stem cells, imposed by the need for a highly dynamic chromatin structure to maintain pluripotency ^50^. Irrespective of the purpose of remodeling, nucleosome turnover necessitates sustained SETDB1 methyltransferase activity for maintaining high H3K9me3 levels at ERVs. This is because newly synthesized H3.3 is devoid of H3K9 methylation, and SETDB1 enzyme is known to establish H3K9me3 only on nucleosomal substrates. Histone H3K9me3 levels decrease in the absence of H3.3 ^21^, showing that histone H3.3 incorporation promotes rather than counteracts establishment of H3K9me3 and the formation of interstitial heterochromatin in mESC.

Collectively, our analyses shed first light into a dynamic heterochromatin state that defies the stereotypical distinction of euchromatin and heterochromatin. While the purpose and mechanism remain only poorly understood, our study provides a novel hypothesis and functional insights for future mechanistic studies.

## ACKNOWLEDGEMENTS

Bioinformatics analyses were performed on resources provided by the Swedish National Infrastructure for Computing (SNIC) at Uppmax server (projects SNIC 2020/15-9, SNIC 2020/6-3, uppstore2018208, SNIC 2018/3-669, sllstore2017057, SNIC 2017/1-508).

## Methods

### Datasets

A list of all used datasets is provided as Supplementary Data Table 1.

### Primary data analysis pipeline

Original FASTQ files were downloaded from GEO and European Nucleotide Archive (ENA) databases (see Extended Data Table 1 for a list of accession IDs). Depending on the deposited source file format, paired-end or single-end reads were mapped to mm9 reference genome using bowtie2 (v 2.3.5.1) ^53^, deduplicated using Picard MarkDuplicates with default settings and filtered for blacklisted regions. Bigwig files were generated from resulting deduplicated and filtered bam files using deepTools (v 3.1.0) ^54^, normalized to 1x Genome Coverage (Reads Per Genomic Content) using --normalizeUsing RPGC --effectiveGenomeSize 2150570000. In 1x Genome Coverage tracks, values below 1 represent a depletion below the genome average whereas values above 1 indicate fold-enrichment above genome average. Unless stated otherwise, downstream analysis was performed on the normalized bigwig files. Where available, we evaluated all experimental replicates (see Extended Data Figure 5). For simplicity, figures show the first replicate.

### Track visualization

We visualize genomic tracks from 1x normalized BigWig files using Integrated Genome Viewer (IGV) ^55^. Y axis corresponds to Reads Per Genomic Content (RPGC).

### Peak calling

Peaks were calculated using MACS2 (v 2.1.2) ^56^ callpeak using parameters: broadPeak, default --broad-cutoff 0.05, corresponding input alignment as control. The three replicates were combined to obtain a reliable set of peaks. A peak is considered if it is called in at least two out of the three replicates. This was performed for both ESC and NSC at initial timepoint (0h). The region considered as a reliable peak is then the result of merging the overlapping called peak regions.

### Bin-based analyses

For the genome-wide coverage analyses in 5kb bins, mean coverage bin files were generated with deepTools multiBigWigSummary bins tool from the final bigwig files using a bin size of 5kb. Where available, input datasets were used to normalize the ChIP-Seq coverage by calculating the log2 of the ChIP/input ratio.

### Repeat element annotation

To filter the many instances of short and fragmented repeat occurrences in the genome, we curated repeat elements from UCSC RepeatMasker as follows: mm9 RepeatMasker track was clustered to merge LTRs and internal sequence into an single continuous interval. For this, adjacent elements (allowing for no gaps) were clustered into single elements, keeping the name of the largest fragment for the cluster (typically the internal sequence, e.g. IAPEz-int, that defines the repeat family). Any single or clustered element shorter than 2kb was removed, clusters larger than 2kb were retained as continuous intervals and used as repeat annotation.

### Curated set of shared IAPEz elements across datasets from different mouse strains

A mm9 reference IAP ERV annotation was obtained from UCSC RepMasker track, merging internal IAPEz-int and flanking LTR intervals and retaining elements longer than 2kb as described above (resulting in 3046 full-length and truncated IAP ERVs). In each available paired-end ChIP-Seq and/or Input dataset, read pairs overlapping with the reference IAP ERVs were retained if at least one of the two reads was unambiguously assigned to one genomic match (thus, the pair could be niquely mapped with bowtie2 mapping quality MQ > 10). One or more such anchoring read was used as evidence that the IAP ERV insertion is present in the respective mouse strain. Evidence from all ChIP-Seq and matched inputs were collectively used to define the IAP ERVs present in a given cell line/mouse background. Finally, the annotation of 2640 shared IAP ERVs was obtained by retaining only those IAP ERVs present in all the cell lines (see Extended Data Figure 10).

### Annotated bins

Genome-wide 5kb bins were intersected with our repeat annotation using bedtools (v2.27.1) intersect ^57^. Bins were assigned the repetitive element group of the element that overlapped the most with each bin, in the unusual case where the same bin overlaps more than one repetitive element.

### Annotated peaks

A peak list from endogenous H3.3 ChIP-seq ^21^ was intersected with peaks called from ^9^ as described above. Any amount of overlap > 0 was considered. Other loci of interest were also intersected with these peaks in the same fashion: IAPEz and ChromHMM15 regions. Since ChromHMM15 is a partition of the genome, a peak is expected to fall in at least one category. For the cases where a peak overlapped with more than one ChromHMM annotation, the largest overlap was reported.

### Peak density heatmaps

Values were calculated with deepTools (v 3.1.0) computeMatrix using parameters --referencePoint center and flanking regions of 2,500 base pairs (-a 2500 -b 2500). In the cases where a different flanking region was used, such length is indicated in the plot. The resulting matrices were visualized using matplotlib (v 3.1.1) with the same parameter settings as deepTools plotHeatmap. Unless stated otherwise, the leftmost heatmap in combined plot is sorted by mean coverage per locus and sorting order is applied to the rest of heatmaps in the same row. Profile plots on top of each heatmap are calculated as the mean values of the columns in the matrices generated by deepTools.

### IAPEz flanking H3.3 density heatmaps

Values were calculated with deepTools. In this case the execution mode was reference-point with --regionBodyLength set to 6500. A set of uniquely-mapped BigWig files was generated from the original aligmnent files using deepTools bamCoverage tool with parameters --minMappingQuality 10 --extendReads 150. For paired-end data, coverage was extended to match fragment length.

### Mean read density heatmaps

Mean coverage values over each of the ChromHMM15 regions were calculated with R/rtracklayer library and normalized to their corresponding input value (log2 ratio). Values obtained were visualized using R package pheatmap.

### Statistics

Plot statistics were calculated using R package ggpubr. Size effect estimations were calculated using R package effsize.

### Genome-wide enrichment over 5kb bins for ES and NS cells, timepoints 0h, 3h, 6h and 12h

Bins were generated using deepTools multiBigWigSummary (see Bins). Each bin value was divided by its corresponding input value. Bins with zero values were excluded from this analysis. Additionally, outliers were removed using Tukey’s 1.5 IQR rule for the violin plots.

### Venn and Euler diagrams

Venn diagrams were calculated and visualized using Intervene ^58^. Proportional representations of these as Euler diagrams were visualized with R package eulerr.

### Profile plots

Average plots over genomic regions were generated with ngsplot ^59^, including a modification to produce plots normalized to 1x Genome Coverage. Since ngsplot considers the coverage of actual fragments defined by paired-end reads for more realistic occupancy profiles, the y axis reports average Fragments Per Genome Coverage (FPKM) rather than RPKM.

---

**Extended Data Figure 1 - related to Figure 1a, b.**
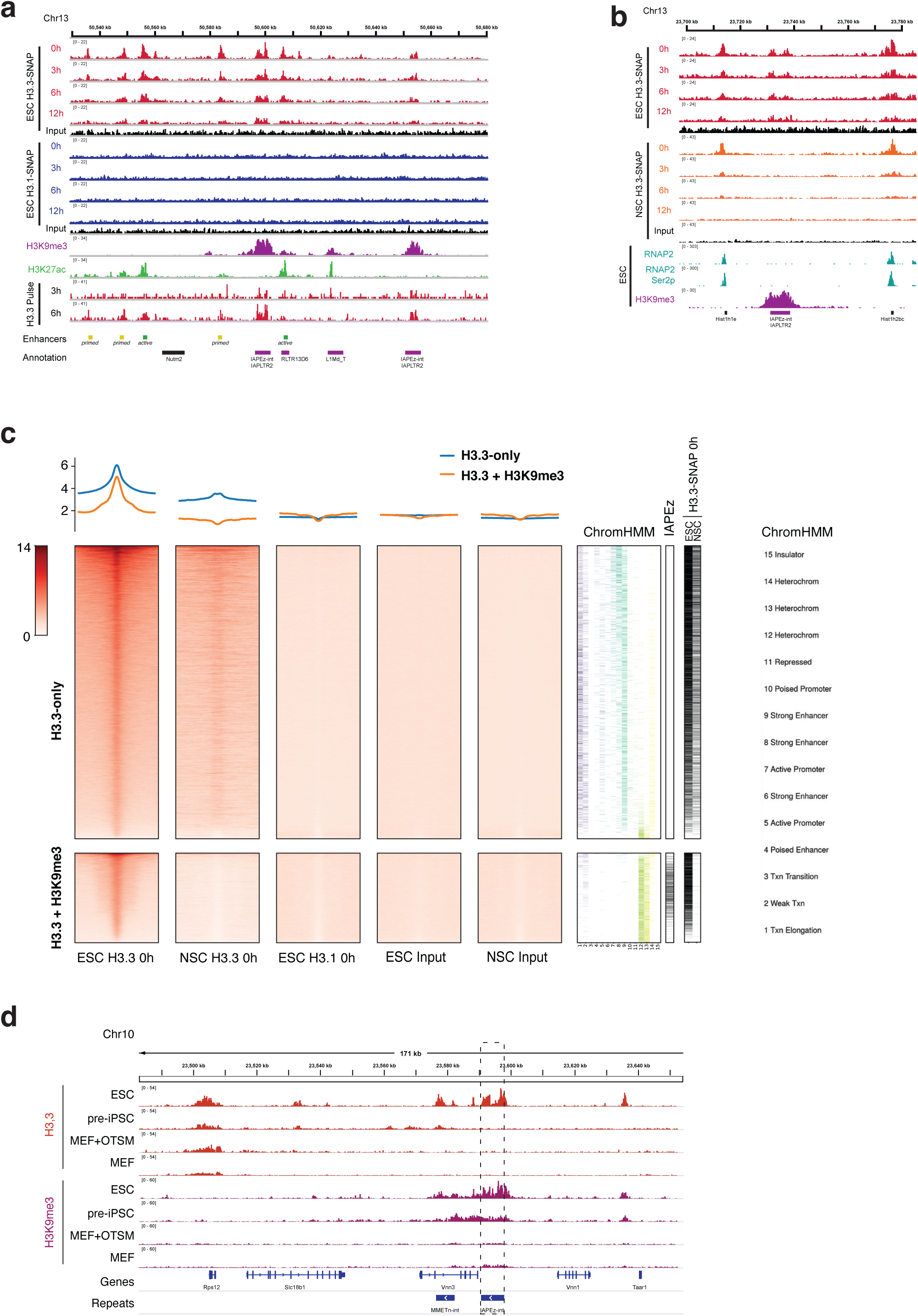
Additional examples and controls of H3.3 enrichment at active regions and interstitial heterochromatin. **a**Genome tracks example of time-ChIP data (H3.3-SNAP and H3.1-SNAP timepoints 0h throughout 12h) in mouse embryonic stem cells (ESC), together with H3.3-HA pulse data ^9,34^, H3K27ac and H3K9me3 ^20,51^. H3.3-SNAP and H3K27ac enriched regions show overlap with active and primed enhancers. H3.3-SNAP and H3K9me3 coincide with IAP ERV (see also genome-wide analysis in Extended Data Figure 3). H3.1-SNAP and Input controls show no particular enrichment. All tracks were normalized to Reads Per Genomic Content (RPGC). **b** Genome tracks example showing time-ChIP H3.3 data in both mouse embryonic stem cells (ESC) and neural stem cells (NSC), together with H3K9me3 ChIP ^51^ and RNA Polymerase II ChIP ^61^ at Hist1 histone gene cluster. H3.3 enrichment and turnover at IAP elements is observed in ESC but not NSC, whereas H3.3 enrichment and turnover at the housekeeping genes is maintained in NSC. **c** Density heatmaps and average profiles (RPGC) of time-ChIP H3.3-SNAP in embryonic stem cells (ESC) and neural stem cells (NSC) and H3.1-SNAP at timepoint 0h in ESC over endogenous H3.3 peaks in ESC coinciding (H3.3+H3K9me3) or not (H3.3-only) with H3K9me3 peaks ^21^. Overlap with 15-State ChromHMM annotation ^37^ states, IAP ERV elements (IAPEz-int and flanking LTRs from mm9 RepeatMasker track), and H3.3-SNAP peaks called in ESC and NSC is shown. Additionally, ESC and NSC input data is also shown. **d** H3.3 and H3K9me3 profiles along a reprogramming time-course from mouse embryonic fibroblasts (MEF) ^35^. Early reprogramming intermediate (48h after expression of reprogramming factors Oct4, Nanog, Klf4, Myc) and late reprogramming intermediate stable cell line (pre-iPSC) are compared to MEF and ESC. All tracks are plotted on the same scale as RPGC. H3K9me3 is first enriched at IAP ERV at the pre-iPSC intermediate and further increases in ESC. H3.3 is exclusively acquired at IAP ERVs in ESC, while nearby housekeeping gene *Rps12* is enriched consistently in all cell types.

**Extended Data Figure 2 - related to Figure 1d.**
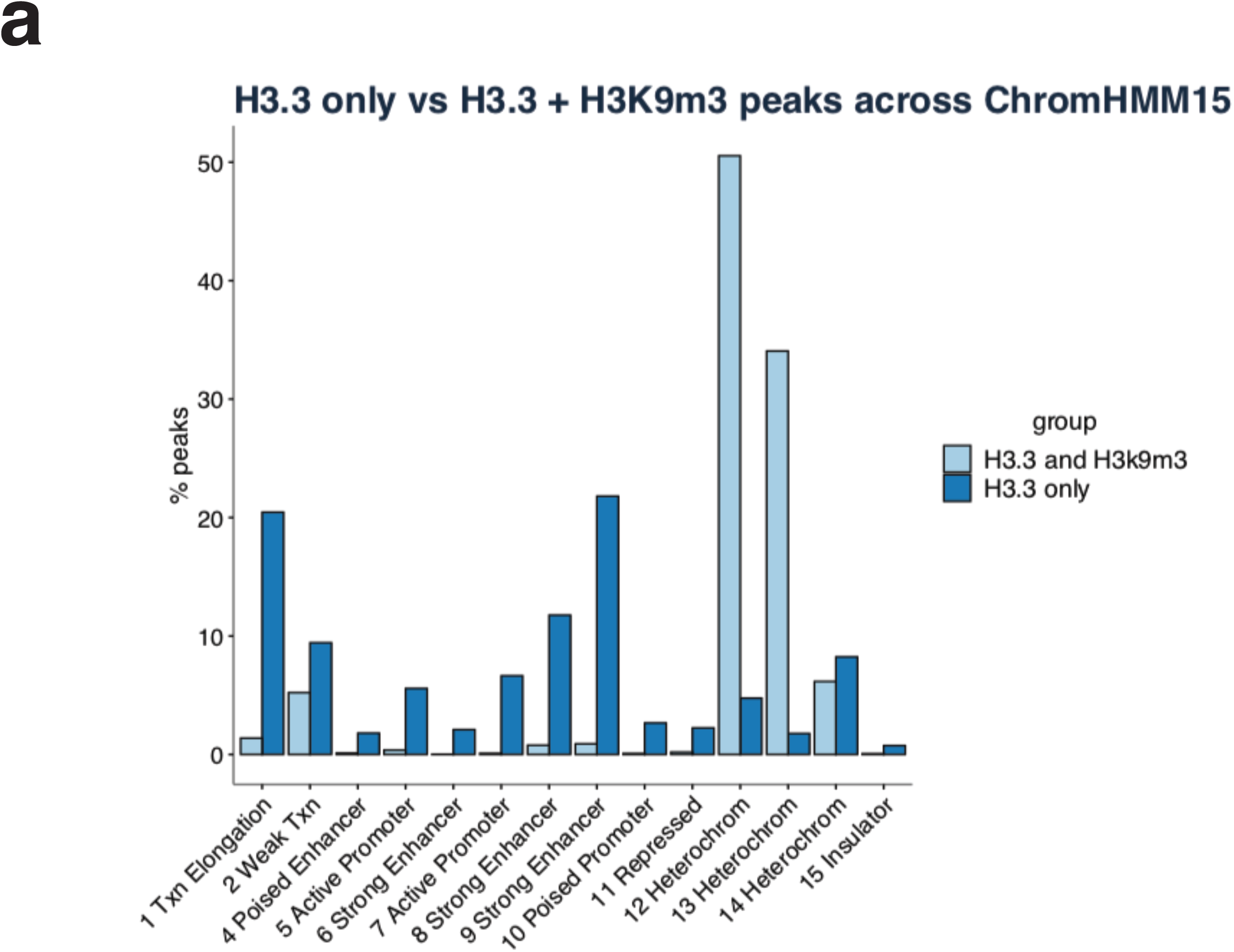
Annotation of H3.3 peaks using 15-State ChromHMM model. Barplot showing relative overlap of “H3.3-only” and “H3.3+H3K9me3” peak sets ^21^ with 15 State ChromHMM annotation, grouped by coincidence with H3K9me3 peaks. Each peak is assigned the ChromHMM15 state that overlaps with it the most. Amounts are shown as a percentage of the total peak set. H3.3-only peaks overlap the most with strong enhancers and transcription elongation states, whereas H3.3+H3K9me3 peaks overlap with heterochromatin states 12-14, defined by absence of H3K4me3, H3K4me1, H3K27ac, H3K36me3 and RNAP2 marks.

**Extended Data Figure 3 - related to Figure 1e.**
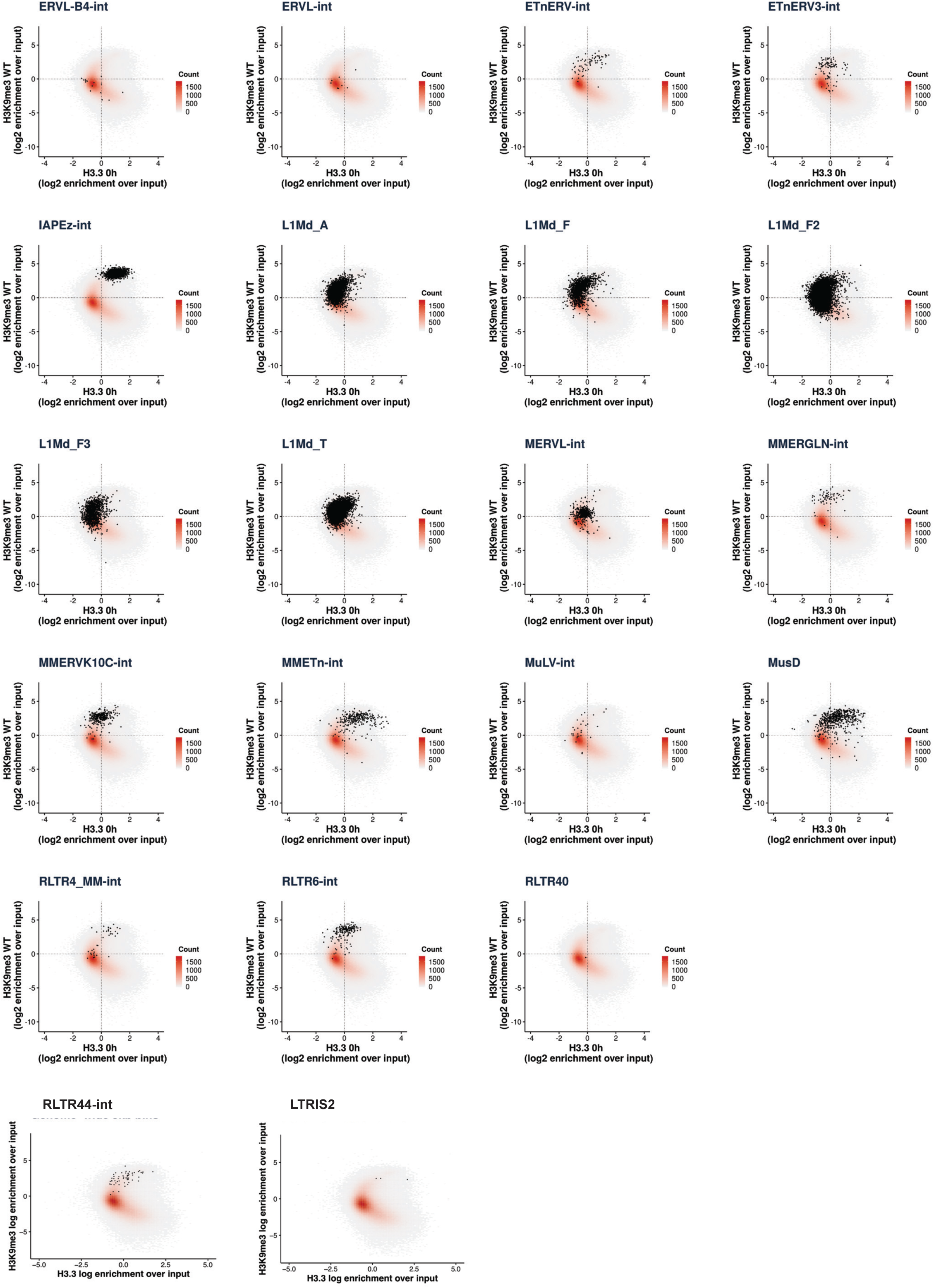
Genome-wide correlation of H3.3 and H3K9me3 is shaped by a subset of ERV families. Scatter plot showing relationship between H3.3 and H3K9me3 signal assessed over 5kb bins genome-wide. Genome was partitioned in 5kb bins and coverage enrichment was calculated as log2 fold change of each signal over their corresponding input. Color of each tile represents point density of all the bins with similar H3K9me3 and H3.3 values (on a tile-resolution of 0.1). RepMasker annotation was intersected with 5kb bins for major repeat families. Bins that overlap with the corresponding repeat type are shown in black. In the rare case where more than one element overlapped with the same bin, the repeat type that overlapped the most was adjudicated.

**Extended Data Figure 4 - related to Figure 1e.**
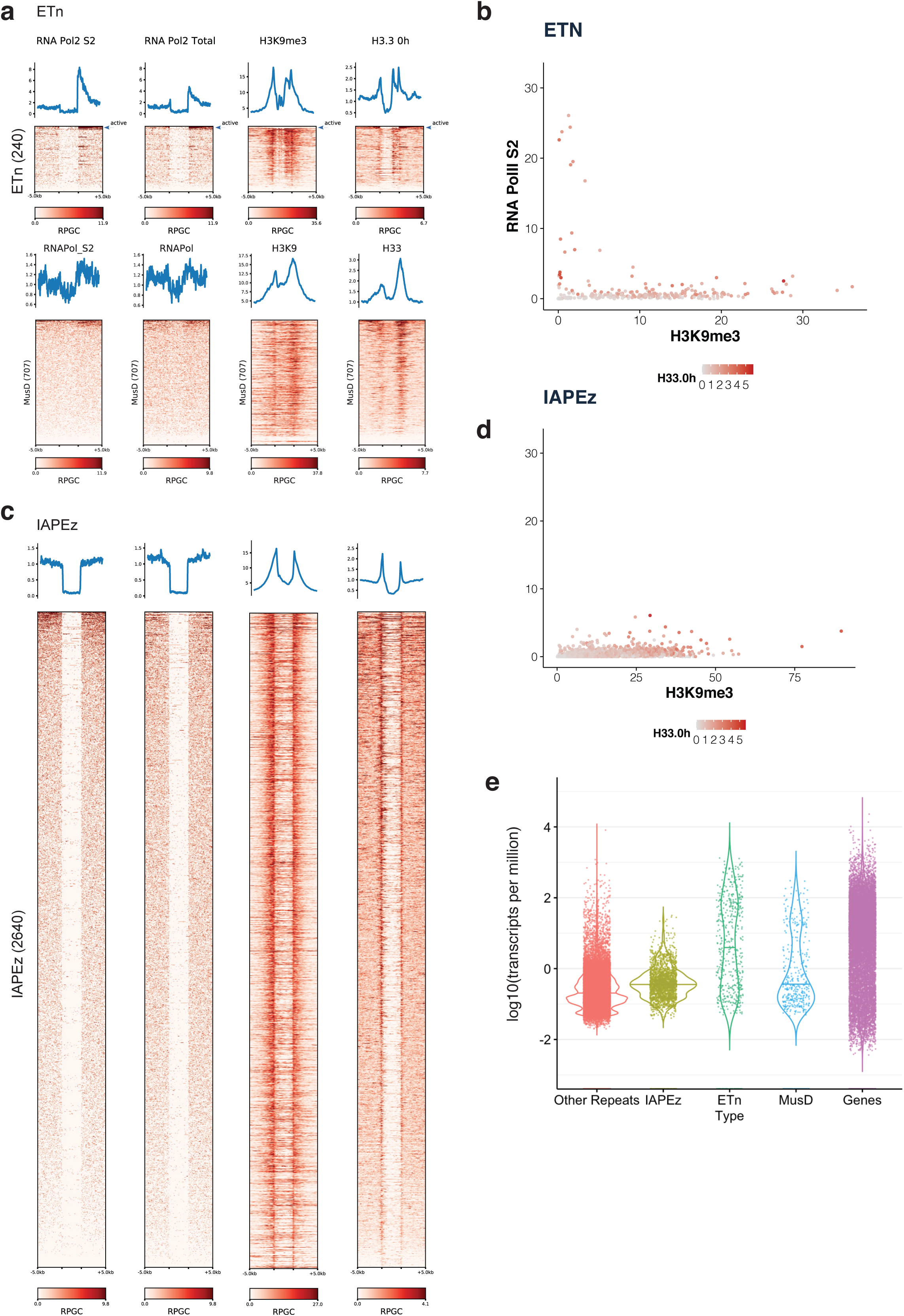
ETn but not IAP ERV families contain a small number of active elements in the annotated genome. **a**Heatmap showing RNA Polymerase II and H3K9me3 signal ^51,61^ over unique 5kb flanking regions of ETn and MusD repeat elements. Among the annotated ETn elements, a small subset (marked with a blue arrow) show high RNA Polymerase II signal at their 3’ flanking region together with absence of H3K9me3, which suggests that these individual elements are transcribed at high levels, whereas the remaining elements are in a repressed or inactive state. **b** Scatterplot showing enrichment of RNA Pol II CTD Serine 2 phosphorylation versus H3K9me3 ^51,61^ over ETn elements and 1kb flanking regions, using only uniquely mappable reads. H3.3 enrichment is overlaid as colorscale. ∼10 elements appear actively transcribed and lack H3K9me3, enrichment of histone H3.3 in these cases may stem from transcription associated nucleosome turnover. **c** Heatmap showing RNA Polymerase II and H3K9me3 signal over 5kb flanking regions of annotated IAP elements. No actively transcribed IAP ERVs are discernable by RNA Polymerase II signal. **d** Scatterplot showing enrichment of RNA Pol II CTD Serine 2 phosphorylation versus H3K9me3 ^51,61^ over IAP elements and 1kb flanking regions, using only uniquely mappable reads. H3.3 enrichment is overlaid as colorscale. **e** Violin plot of RNA-Seq transcript levels for genes and selected repeat elements. RNA-Seq read counts were mapped to mm9 and read counts were extracted over RefSeq genes and RepMasker annotation. Read counts were normalized to the total base pairs covered by each gene or repeat instance (RPK, reads per kilobase) and scaled to transcripts per kilobase million (TPM) and plotted as in log scale.

**Extended Data Figure 5 - related to Figure 2a-e.**
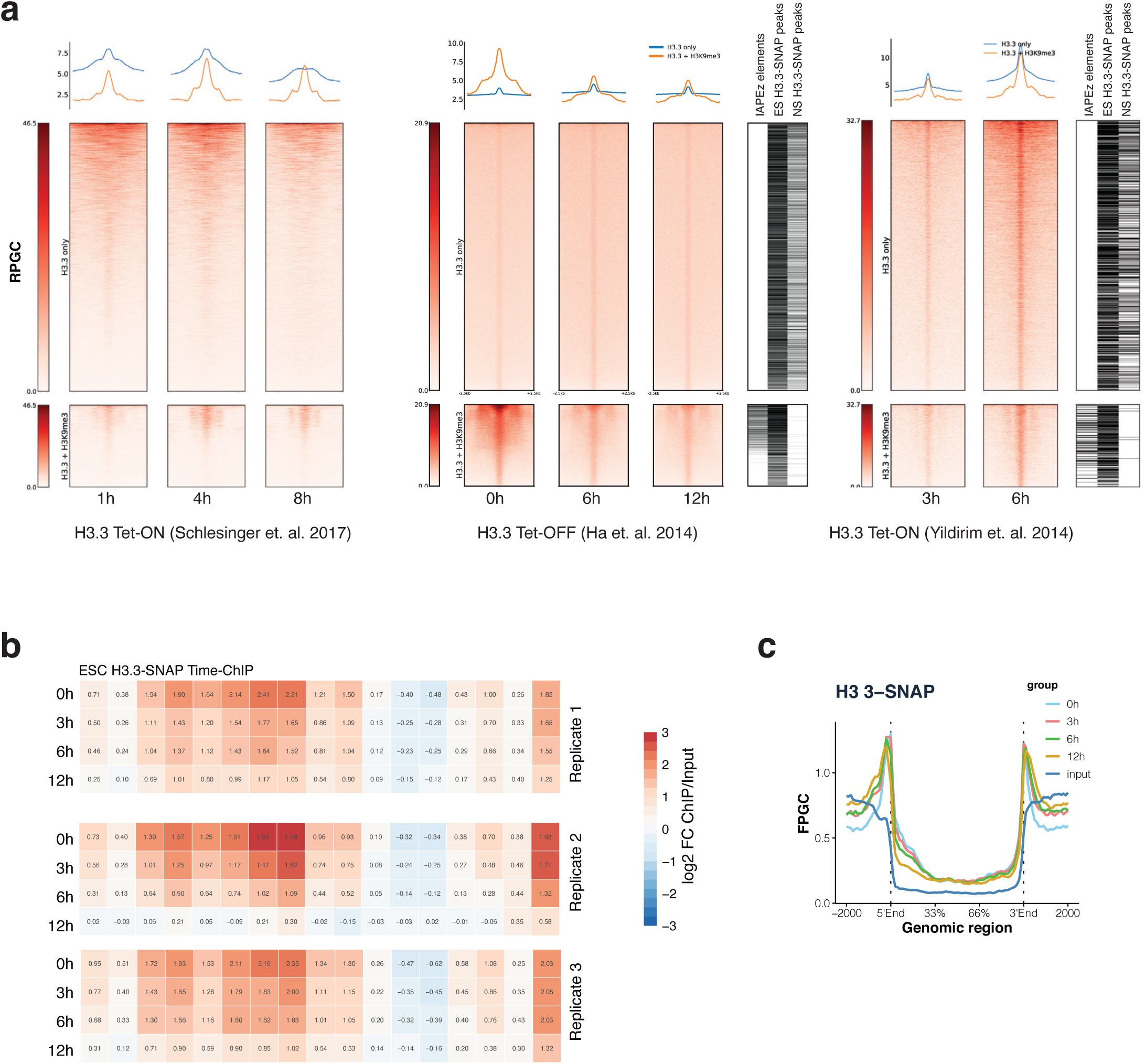
Analysis of additional H3.3 pulsing datasets. **a**Peak Density Heatmaps and average profiles of H3.3 pulse or chase data from three different studies in mouse ESC ^5,8,34^ at “H3.3-only” and “H3.3+H3K9me3” peak sets ^21^. **b** Mean read density heatmap showing enrichments of H3.3-SNAP time-ChIP (log2 fold-change over input) over 15 ChromHMM states as well as H3K9me3 and H3.3+H3K9me3 enriched regions ^21^. Same as Figure 2d but with additional replicates ^9^. **c** Average coverage of H3.3-SNAP time-ChIP uniquely mappable reads over 2640 shared IAP ERVs. Fragments defined by paired-end reads were piled up and normalized by 1x Genome coverage (Fragment Per Genomic Content, FPGC)

**Extended Data Figure 6 - related to Figure 2d.**
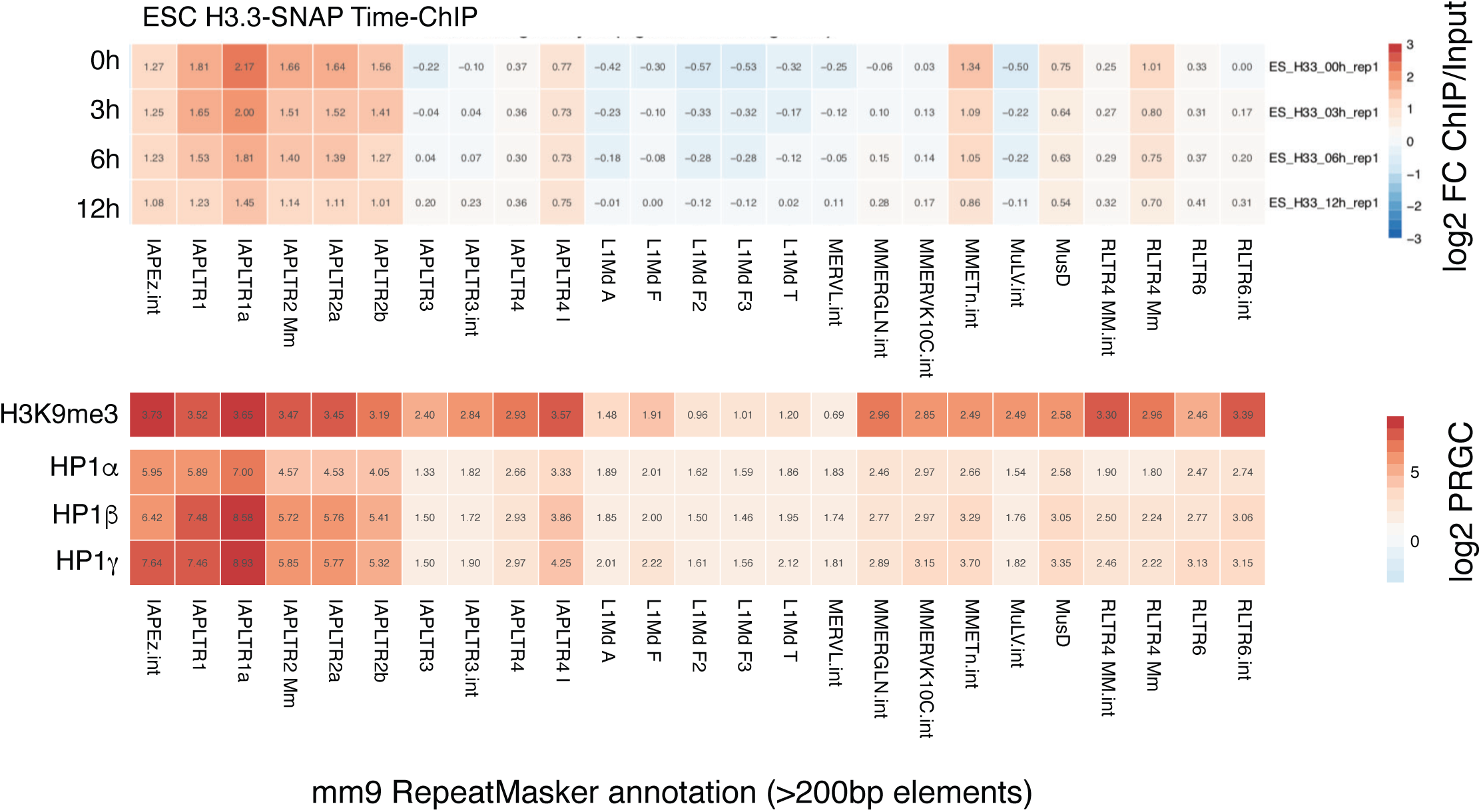
Analysis of repetitive element families. **a**Mean read density heatmaps of time-ChIP H3.3-SNAP data at a major repetitive elements from RepeatMasker annotation (filtered for elements larger than 200bp). Each category enrichment is calculated as log2 fold change of mean coverage over input. IAPLTR1a shows the highest enrichment at time point 0h and turnover. Most IAP LTRs and internal sequences, as well as ETn and MusD elements show both enrichment and H3.3 turnover.

**Extended Data Figure 7 - related to Figure 3a, d.**
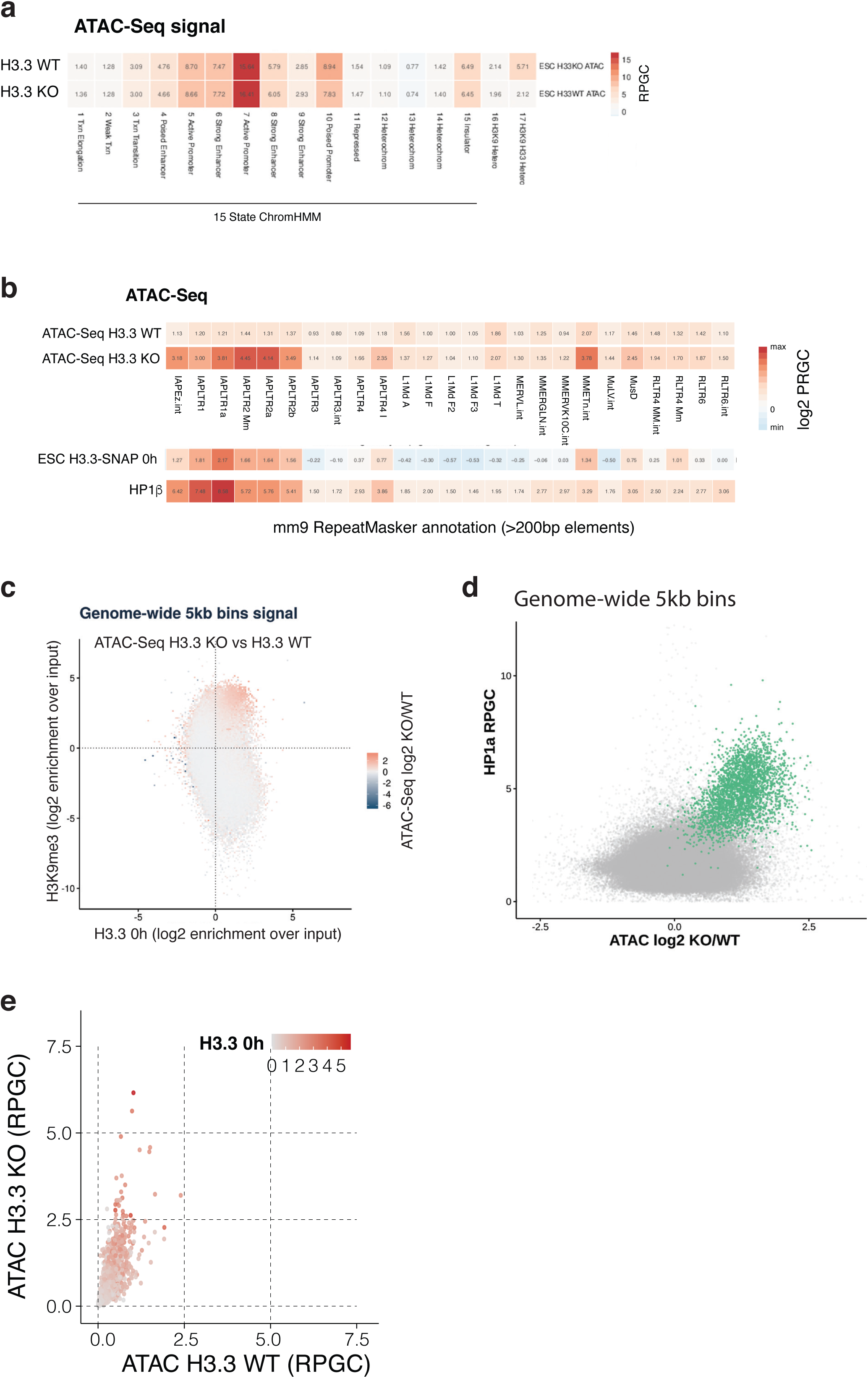
Genome-wide maintenance of accessibility in the absence of H3.3KO cells independent of active or inactive chromatin state, and selective gain in accessibility at H3K9me3-H3.3 co-enriched regions. **a**Mean read density heatmap of ATAC-seq data of H3.3 wildtype and knockout cells ^20^. Both WT and KO show very similar genome-wide read density levels across all 15 ChromHMM States. While H3K9me3-only regions remain inaccessible, a striking ∼3-fold increase in ATAC-Seq signal is found at H3.3 + H3K9me3 regions in H3.3KO cells. **b** Mean read density heatmap of ATAC-seq data of H3.3 wildtype and knockout by repeat family. Increase in accessibility is observed for those repeat families that are enriched in H3.3 in wildtype cells. **c** Tile-plot showing relationship between H3.3 and H3K9me3 signal assessed over 5kb bins genome-wide. Each tile color shows log2 fold enrichment of average ATAC-seq signal for H3.3 KO over WT of all the bins with similar H3K9me3 and H3.3 values (on a tile-resolution of 0.1). A majority of zero values show ATAC-seq signal is very similar genome-wide, with H3.3 KO showing highest enrichment over WT on regions both H3.3- and H3K9me3-enriched. **d** Scatter plot of 5kb bins, showing HP1a occupancy ^52^ versus log2-fold change in ATAC-Seq signal upon H3.3 knockout. Bins overlapping with IAP ERVs are drawn in green. **e** Scatter plot showing uniquely mappable ATAC-Seq reads (RPGC) in H3.3 KO versus WT cell lines, at 2640 shared IAP ERVs and 1kb flanking regions. H3.3-SNAP 0h density by unique reads is overlaid as color scale.

**Extended Data Figure 8 - related to Figure 3a.**
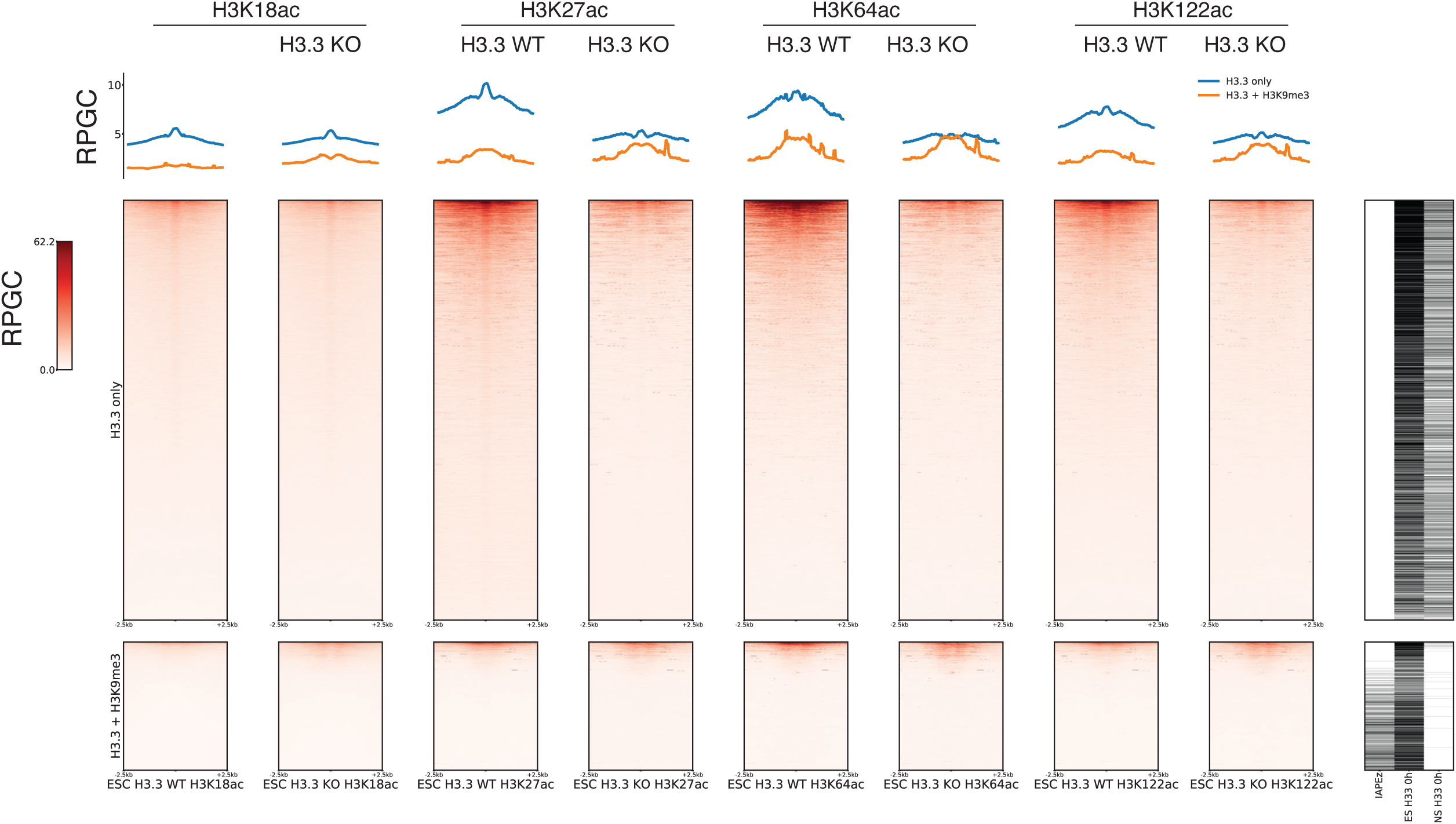
Loss of H3.3 does not lead to an increase in histone acetylations at H3.3+H3K9me3 regions. **a** Density Heatmaps and average profiles of H3K18ac, H3K27ac, H3K64ac and H3K122ac marks for H3.3 WT and KO samples ^20^ at H3.3-only and H3.3+H3K9me3 peak sets. As previously described, histone acetylation, present at a subset of H3.3 peaks, is reduced in the absence of H3.3 ^20^, but basal levels at H3.3+H3K9m3 regions are not changed.

**Extended Data Figure 9 - related to Figure 3f.**
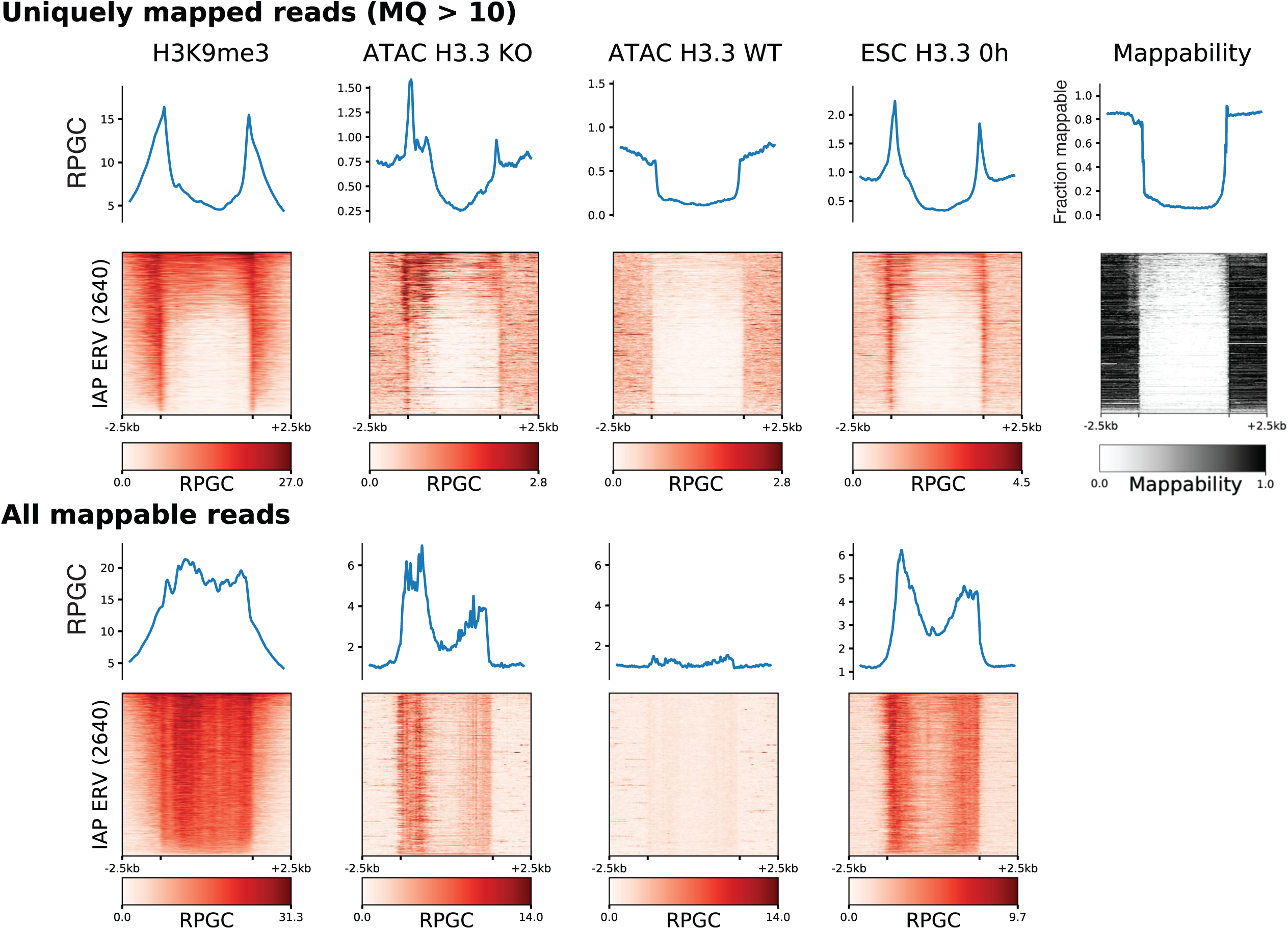
Uniquely mappable reads show widespread increase in DNA accessibility over individual IAP ERV elements. Density heatmap of uniquely mappable paired-end reads within and adjacent to 2640 shared IAP ERVs for H3K9me3, H3.3 KO and WT ATAC-seq, and Time-ChIP H3.3-SNAP at timepoint 0h. Top row shows average profiles. Uniquely mapped reads at flanking regions allow to distinguish among individual instances of repetitive elements, showing that the observed effect is pervasive across such repeat elements. Lower row shows the corresponding unfiltered signal including multi-mappable reads for the same datasets at the same IAP ERVs set. Multimappable reads were assigned randomly to one of the ambiguous matches. On the rightmost column, a 100bp mappability track that flanking regions and some internal sections of IAP ERVs are uniquely mappable.

**Extended Data Figure 10.**
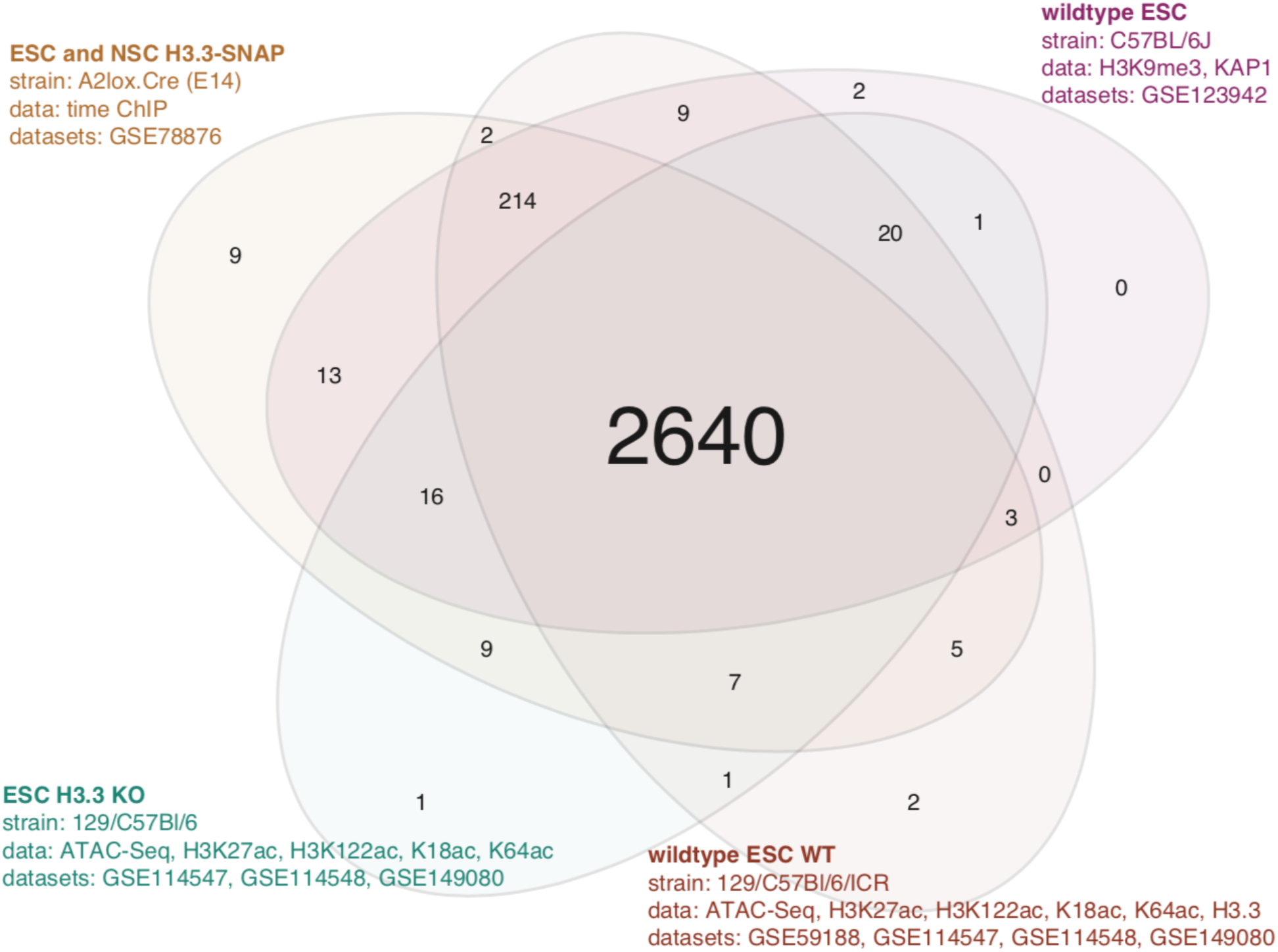
Consensus IAP ERV elements shared across datasets from different mouse strains. Venn diagram showing overlap between the datasets used to collect evidence from shared IAP ERV elements on a per-locus basis. Since evidence was based on paired-end reads anchored in the unique flanking region (see Methods), we could only include paired-end datasets in the census. For each individual study, a IAP ERV element was retained if it showed evidence on any of the datasets. From a total set of 3046 full-length IAP ERVs obtained from UCSC RepMasker track, a curated consensus final dataset of 2640 loci was obtained.

